# Exploring the Regeneration Dynamics and Conservation Threats to Endemic *Rhododendron kesangiae* (D.G. Long & Rushforth) in Bhutan’s National Botanical Park at Lungchutse, Dochula

**DOI:** 10.1101/2024.09.03.610747

**Authors:** Chogyal Tshering Dolkar, Yonten Dorji

## Abstract

*Rhododendron kesangiae* is a species endemic to the Eastern Himalayas named after the Queen mother of Bhutan, HRH Ashi Kesang Choden Wangchuk. The study on the regeneration and conservation threat of *Rhododendron kesangiae* is of utmost importance in this changing climate and growing human interference in the natural population of Rhododendrons. However, there is limited documentation and study on this species’ ecology, regeneration, and conservation status. Therefore, this study was undertaken to assess the regeneration ecology and conservation status of R. *kesangiae* in one of Bhutan’s National Botanical Park at Lungchutse, Dochula. The study area was divided into six transects along the North East and North West aspects. The regeneration status of R. *kesangiae* was determined by counting the number of seedlings, saplings and adults in the 5 × 5 m transects. Composite soil samples were collected from the 5 × 5 m quadrant. Associated tree species data were gathered from 20 × 20 m quadrants in each plot. The study found that the regeneration status of R. *kesangiae* was fair regeneration with the seedlings ≤ saplings > adults. The most dominant tree species found were *Tsuga Dumosa* and the least dominant species was *Juniperus recurva*. Soil parameters such as soil moisture significantly impacted the regeneration of *R. kesangiae* (*r* = 0.52, *p* = 0.003). Precipitation had a significant impact on the regeneration and growth of R. *kesangiae* (*r* = 0.37, *p* = 0.043), while other environmental variables such as slope, temperature and elevation did not show a significant impact. The conservation threats were documented using Miradi. This study sheds important light on the species’ regeneration ecology and conservation status highlighiting the importance of monitoring and conservation efforts to ensue their long-term survival and keeping them from falling into the highest conservation threat categories.

## 1. Introduction

*Rhododendron* originates from the Greek word “rhodon” signifying “rose” and “dendron” signifying “tree” (Paul et al., 2015). *Rhododendron* was first described by Carl Linnaeus in his work “Genera Plantarum” in 1737 (Mao, 1970). *Rhododendron* is considered one of the most vast and diverse plant species, occurring at high elevations and having paramount ecological and economic importance. The genus is considered one of the oldest flowering plants, having lived for around 100 million years in temperate northern hemisphere locations (de Milleville, 2002). *Rhododendron* is an important genus in the Ericaceae family, with approximately thousands of species in Asia, Europe, and North America (Paul et al., 2015).

The Bhutan Himalayas, which include a major part of the Eastern Himalayas, are characterised by their challenging topography and diverse climate. As vegetation remains so diverse, it serves as the foundation for a significant portion of floristic diversity, particularly in *Rhododendrons*, which make up for around 4% of the genus’ entire global population (Namgay & Sridith, 2020). In Bhutan, 46 *Rhododendron* species have been identified and classified into two subgenera: *Rhododendron* and *Hymenanthes* (Namgay & Sridith, 2021).

*Rhododendron kesangiae* belongs to the family *Ericaceae* and is an evergreen shrub endemic to the Eastern Himalayas, where it grows at an altitude of 2600–3400 masl. R. *kesangiae* was first described by Long & Rushforth (1995) in honour of the Queen mother of Bhutan, Ashi Kesang Choden Wangchuck. It is locally known as Tala and is usually found in Bhutan’s fir and hemlock forests (Namgyal & Long, 1995).

There are numerous studies on *Rhododendron* species both inside and outside Bhutan, but limited studies on *Rhododendron kesangiae*, which is a critical species of the sub-alpine and alpine ecosystems. The lack of study on *Rhododendron kesangiae*, along with its restricted range due to overexploitation and its classification as a tertiary relict species (Lhaki, 2021), raises questions about possible loss of biodiversity and increases the species’ conservation threat level. The most delicate environment in the Himalayas is found in the subalpine to alpine transition zone, which includes tree line. There is no doubt that the *Rhododendron* family, the sole plant group with continuity in the ecotone mentioned above, sustains biological sustenance in this sensitive area (Singh et al., 2003). Deforestation and unsustainable extraction by locals for fuel and incense pose the biggest risks to *Rhododendron*s (Singh et al., 2003).

*Rhododendron* restoration and preservation in the wild encourage the occurrence of other elements of biodiversity. It provides food preserve for a variety of birds at an altitude gradient (Singh et al., 2003). Thus, understanding the regeneration dynamics and conservation status of the species is important to help preserve the species in its natural habitat (Tiwari et al., 2018).

Regeneration and conservation threat studies of *Rhododendron kesangiae* have become essential in this changing climate and increased human activity in the natural population of *Rhododendrons* in the Himalayan areas (Singh et al., 2009), which is at significant risk of habitat loss and fragmentation. Monitoring and conservation efforts are critical to ensuring their long-term survival and keeping them from falling into the highest conservation threat categories.

The objectives of this research are to i) assess the current regeneration status of the *Rhododendron kesangiae* population in Lungchu Tse, Dochula, ii) and to identify and evaluate the conservation threats facing this species. iii) Additionally, the study aims to investigate the ecological factors that influence the regeneration and growth of *Rhododendron kesangiae*. To achieve these objectives, the research will address several key questions: What is the current regeneration status of *Rhododendron kesangiae* in Lungchu Tse, Dochula? What are the primary conservation threats to *Rhododendron kesangiae* in this area, and how can these threats be prioritized for effective conservation planning and action? Furthermore, what are the key ecological determinants, such as soil nutrient composition and environmental variables, that affect the regeneration and growth of *Rhododendron kesangiae*? Through this comprehensive approach, the study seeks to contribute valuable insights into the conservation and management of this endemic species, promoting its sustainable future in Bhutan’s unique ecosystem.

## 2. Materials and Method

### 2.1 Study site and experimental design

The study was conducted in the Lungchutse region of Dochula (27.507668°N and 89.75287°E). It is located at an altitude of 3100 to 3569 metres above sea level and covers around three kilometres (Bhutaninbound, 2020). The high altitude of the study area provides a distinctive setting for studying how human activities affect the environment and how plants and animals adapt to high elevations (Singh, 2016).

Recently, human activities such as tourism and garbage dumping have increased in the area, which poses a risk of harming the ecosystem at the Dochula Pass and surrounding areas (thirdeyemom, n.d.).

The Lamperi region and the eastern slope of the Dochula Pass provide a representative sample of the vegetation found throughout the Himalayan range, including the dry western and moist east Himalayas (Wangda and Ohsawa, 2006). Dochula, located at 3185 masl, received an average annual precipitation of 1575.5 mm between 1999 and 2004 (Dorji and Gurung, 2018). In accordance with BLCA (Bhutan Land Cover Assessment 2010), the research region comprises six different types of vegetation: Chirpine forest (CP), Cool Broad-leaved Forest (CBL), Warm Broad-leaved Forest (WBL), Mixed Conifer Forest (MC), Meadows (MD), and Agricultural land (AGR) (Grierson & Long, 1983; Wangda & Ohsawa).

**Figure 1:**
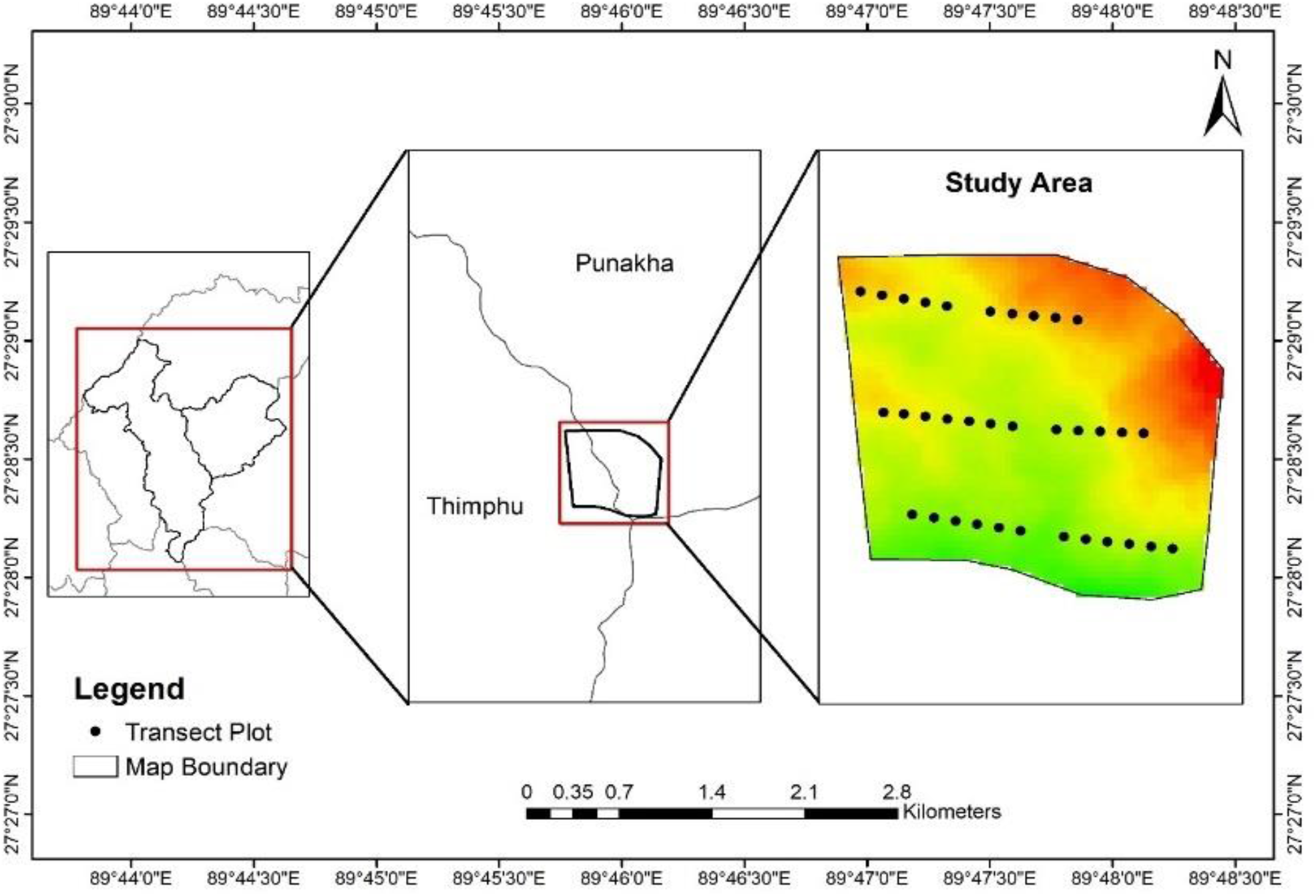
Study area map

### 2.2 Sampling Design

The transect sampling method was used for this study, where six transects (T1-T6) were established at a 500 m interval between two transects. Each transect had five plots, for a total of thirty plots. Tree data was collected using 20 × 20 m quadrants, whereas regeneration was measured using 5 × 5 m inner quadrants separated by 100 m intervals. Transects were laid out on the Northeast and Northwest aspects. Individuals of *Rhododendron kesangiae* were categorized according to their height and basal diameter.

#### 2.2.1 Data collection

Seedlings were defined as those standing less than 20 cm tall. Saplings were defined as individuals that were taller than 20 cm and had a basal diameter less than 4 cm. R. *kesangaie* individuals with a basal diameter greater than 4 cm were classified as adults, and this classification is in line with the established method used by Sharma et al. (2020).

The coordinates, elevation, aspect, and slope of each plot were recorded. Tree species growing in association with R. *kesangiae* were also recorded to understand their interaction.

A composite soil sampling method was used to collect soil samples from each plot. The soil sample was collected from a depth of 0-30 cm using soil auger. Five sub-samples were collected (four from each corner and one from the centre) from each 20 by 20 m plot to form a composite soil sample to measure the soil’s physio-chemical properties (Wangmo et al., 2022).

### 2.3 Materials

The materials required are GPS, compass, clinometer, diameter tape, measuring tape, markers and recording book, camera, and SW map. Altitude was measured using an altimeter, aspect using a compass, coordinates with GPS, and slope with a clinometer. To collect soil samples, the materials used were soil probes/auger and container bags (Wangmo et al., 2022).

### 2.4 Data analysis

*Density of Rhododendron kesangiae (seedlings, saplings, and adults)*

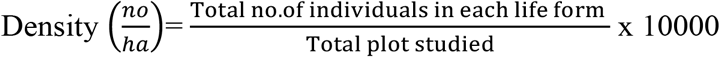

#### Regeneration status

The regeneration structure of the species was determined based on the population size of seedlings and saplings (Lhaki, 2021).

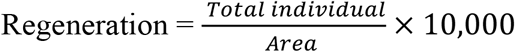

#### Associated tree species

For associated species, correlation was determined to understand the ecological factors affecting the growth of the species and the ecological interaction between the species and associated species. To calculate the most dominant associated species, Important Value Index (IVI) analysis was determined using (Chauhan et al.,2017):

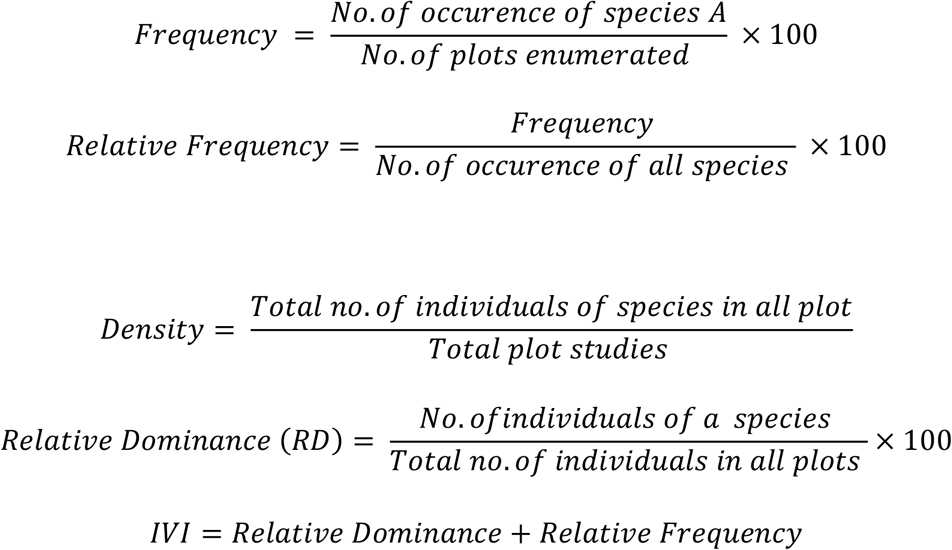

Excel was used to clean, sort, and arrange data for analysis. The arranged data from Excel were imported and analysed with SPSS version 26 to fulfil the research objectives of this study. We had descriptive statistics, graphs, charts, and other statistical tests included. PCORD was used to perform CCA analysis to determine the environmental parameters that influence *Rhododendron kesangiae* growth and regeneration. Miradi was used to record and document the conservation threats to R. *kesangiae* in the study area.

## 3. Results and Discussion

### 3.1 Seedling, Sapling, and Adult Density of *Rhododendron Kesangiae*

To determine the distribution of individuals of R. *kesangiae*, descriptive statistics were calculated. The analysis was conducted on a sample size of 30 plots were sampled with a total of 454 individuals recorded. The mean number of seedlings was 4.70 (*M* = 2, *SD* = 6.39), with the individual range from 1 to 21. Transect 5 had the highest number of seedlings (10.80, *SD* = 7.759), whereas transect 2 had the lowest (0.80, *SD* = 1.095).

The mean number of saplings was 6.13 (*M* = 4.50, *SD* = 6.58), with the individual range from 0 to 32. The highest mean of saplings was recorded in transect 1 (9.00, *SD* = 13.304) and lowest in transect 2 (2.80, *SD* = 2.950). The mean number of adults was 4.30 (*M* = 3, *SD* = 3.07), with the individual range from 0 to 12. The mean number of adults in transect 5 was the highest (9.40, *SD* = 1.673), while transect 2 had the lowest (2.80, *SD* = 2.280), respectively.

**Table 1:** Descriptive Statistics for the Regeneration Status of *Rhododendron kesangiae*.

|  | N | Mean | Median | Mode | SD | Min | Max |
| --- | --- | --- | --- | --- | --- | --- | --- |
| No of seedlings | 30 | 4.70 | 2 | 2 | 6.390 | 0 | 21 |
| No of saplings | 30 | 6.13 | 4.50 | 0 | 6.585 | 0 | 32 |
| No of adults | 30 | 4.30 | 3 | 3 | 3.007 | 0 | 12 |
| Total | 30 | 15.13 | 11 | 11 | 10.45 | 0 | 36 |

Note: N = sample size, SD = standard deviation

### 3.2 Regeneration status of *Rhododendron kesangiae*

The regeneration status of *Rhododendron kesangiae* was determined based on the number of individuals of seedlings and saplings based on the criteria that regeneration is good if seedlings > saplings > adults, regeneration is fair if seedlings > or seedlings ≤ saplings ≤ adults or if seedlings ≤ saplings > adults or if seedlings ≥ saplings and the species have no adults and regeneration is poor if there is only sapling stage of the species even though saplings may be <, >, or = adults. It will be considered as a new regeneration if the species survives only in the seedling stage.

**Figure 2:**
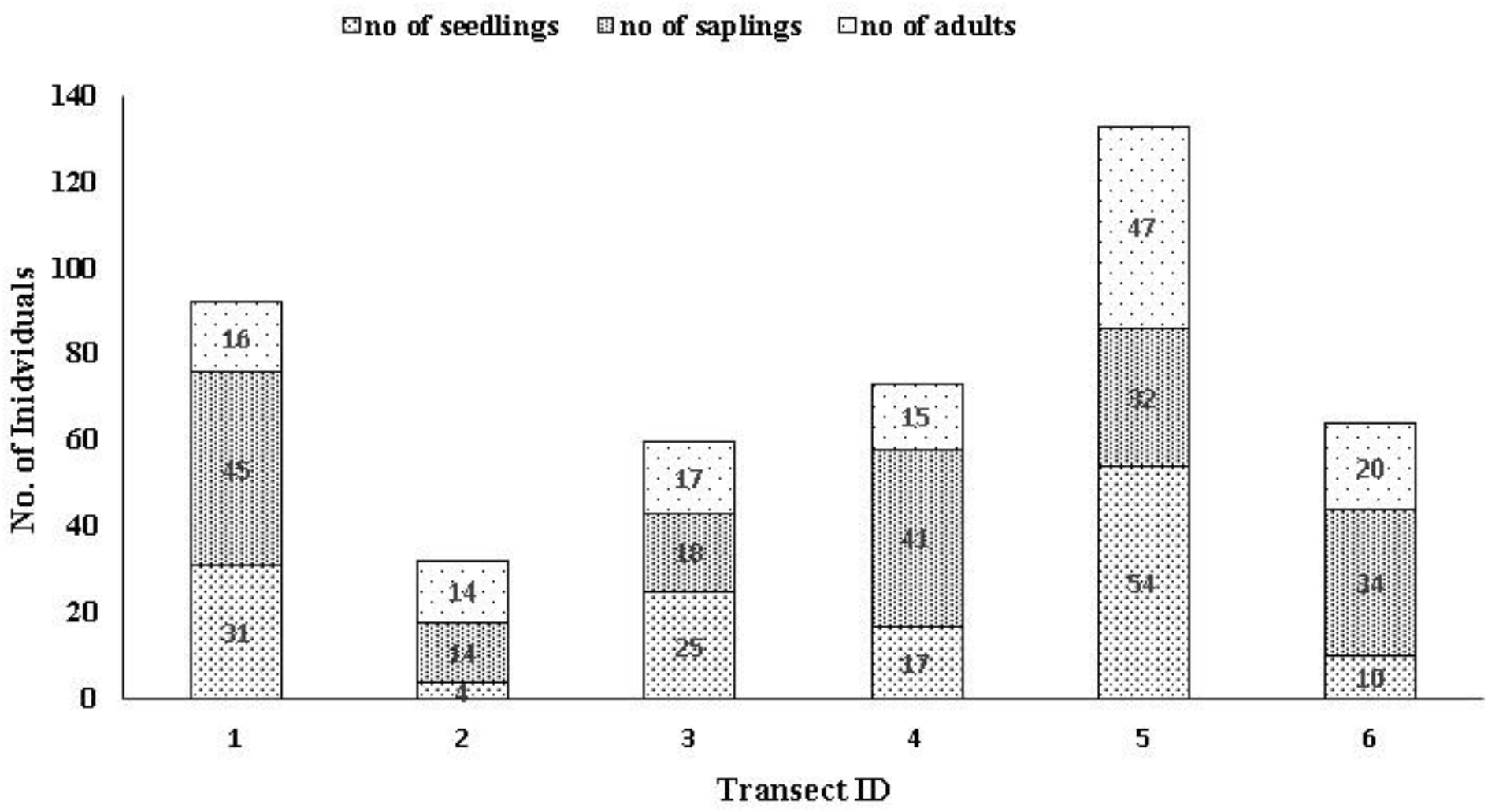
Population structure of *Rhododendron kesangiae*

The analysis of the population structure of *Rhododendron kesangiae*, based on the number of seedlings, saplings, and adults, provides insights into the current regeneration status of the species in Lungchu Tse, Dochula. The study found that the status of regeneration in the study area is fair, with the number of seedlings being more significant than the number of adults. However, the number of saplings is greater than the number of seedlings (141 seedlings ≤ 184 saplings > 129 adults).

The greater quantity of seedlings compared to adults indicates a favourable outlook for regeneration potential, signifying continuous growth and rejuvenation within the population. Similar results were found in a study conducted to investigate the population structure and regeneration status of *Rhododendron* tree species in temperate mixed broad-leaved forests in Tawang and West Kameng districts of western Arunachal Pradesh, India, with 77% of the species exhibiting fair regeneration (Paul et al., 2019).

### 3.3 Regeneration between different aspects

An independent sample t-test was conducted to compare the regeneration between two different aspects (NE and NW), respectively. The analysis revealed that there was no significant difference in regeneration of seedlings (*t* (28) = -0.59, *p* = .558) and saplings (*t* (28) = -0.82, *p* = .415) between the NE and NW aspect of the study area.

**Table 2:** Independent sample t-test between regeneration in different aspects.

|  |  | Levene's Test<br>for Equality of<br>Variances |  | t-test for Equality of Means |  |  |  |  |  |  |
| --- | --- | --- | --- | --- | --- | --- | --- | --- | --- | --- |
|  |  | F | Sig. | t | df | Sig.<br>(2-tailed) | Mean | Std.<br>Error | 95% (CI) |  |
| Regeneration | Equal<br>variances<br>assumed | 2.683 | **0.113** | **-1.059** | **28** | **0.299** | -2.46 | 2.32 | -7.23 | 2.3 |
|  | Equal<br>variances<br>not<br>assumed |  |  | -1.059 | 25 | 0.3 | -2.46 | 2.32 | -7.26 | 2.32 |

These findings suggest that aspect is not a significant factor affecting regeneration in the study area, whereas similar results were obtained in a study conducted by Sharma et al., 2020. The no difference between the aspects can be due to a number of factors, such as uniform regeneration patterns and similarity in microclimatic variability in the study area.

**Figure 3:**
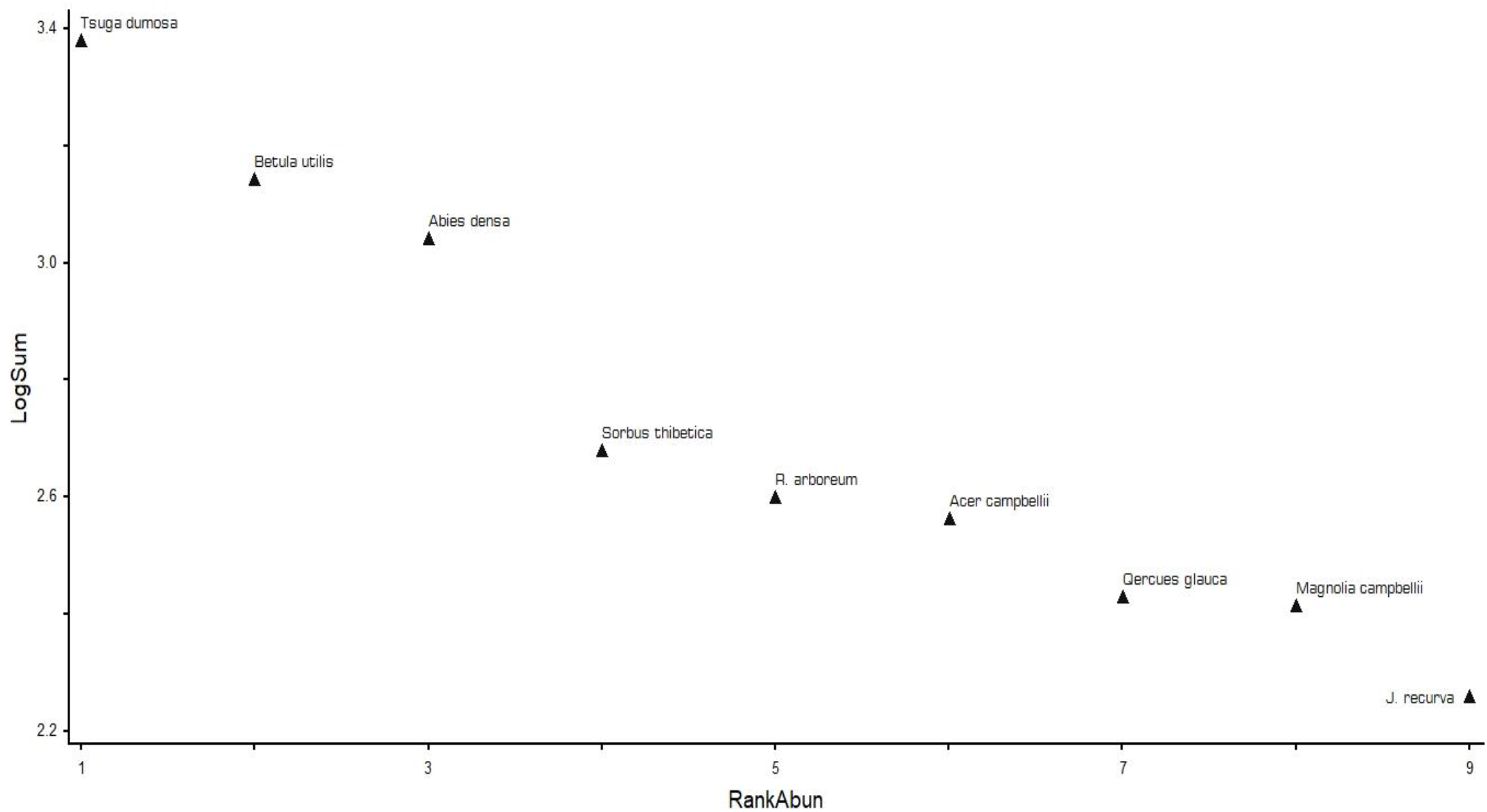
Associated tree species dominance curve

### 3.4 Associated tree species dominance

A total of nine tree species under eight families was recorded across the study site. Family *Pinaceae* had the highest species count (2), followed by a species each in the family *Ericaceae, Fagaceae, Sapindaceae, Cupressaceae, Magnoliaceae, Rosaceae*, and *Betulaceae. Tsuga Dumosa* was also recorded as the most dominant tree species in IVI calculation with an IVI of 99.60, followed by *Betula utilis* and *Abies densa* with an IVI of 62.09 and 40, respectively. The least dominant species was *Juniperus recurva*, with an IVI of 4.38. The community structure and composition of an area can be understood by studying the associated species. R. *kesangiae* is found to be growing in association with Hemlock, Fir, and other Rhododendron species (Pual et a., 2001). Identifying the dominant tree species, such as *Tsuga dumosa, Betula utilis*, and *Abies densa*, might help establish their potential impact on *Rhododendron kesangiae* survival and regeneration.

### 3.5 Relation between associated tree species and soil parameters

We gain valuable insights into what influences *Rhododendron kesangiae’s* growth and regeneration by analysing the relationships between temperature, precipitation, altitude, slope, species composition, and various soil factors (such as organic carbon, nitrogen, organic matter, moisture, phosphorus, potassium, pH, and electrical conductivity). While soil organic carbon, nitrogen, and organic matter exhibited relatively moderate associations, soil moisture demonstrated a significantly stronger association with species composition (*r*= 0.323 - 0.540, *rs-q* = 0.104 - 0.292, *tau* = 0.172 - 0.329), soil moisture (SM) displayed relatively small associations with species composition (*R* = 0.602, *rs-q* = 0.362, *tau* = 0.415).

The species composition and phosphorus (P) showed relatively small to moderate positive correlations (*r*= 0.103 - 0.381, *rs-q* = 0.011 - 0.145, *tau* = -0.131 - 0.283). The moderately positive correlations for potassium (K) were as follows: *r* = 0.234 - 0.485, *rs-q* = 0.001 - 0.236, *tau* = 0.320 - 0.320. The results for pH were inconsistent, showing strong negative correlations in some instances (*r*= -0.420, *rs-q* = 0.177, *tau* = -0.305) and high positive correlations in others (*r* = 0.105 - 0.629, *rs-q* = 0.011 - 0.396, *tau* = 0.037 - 0.323).

**Figure 4:**
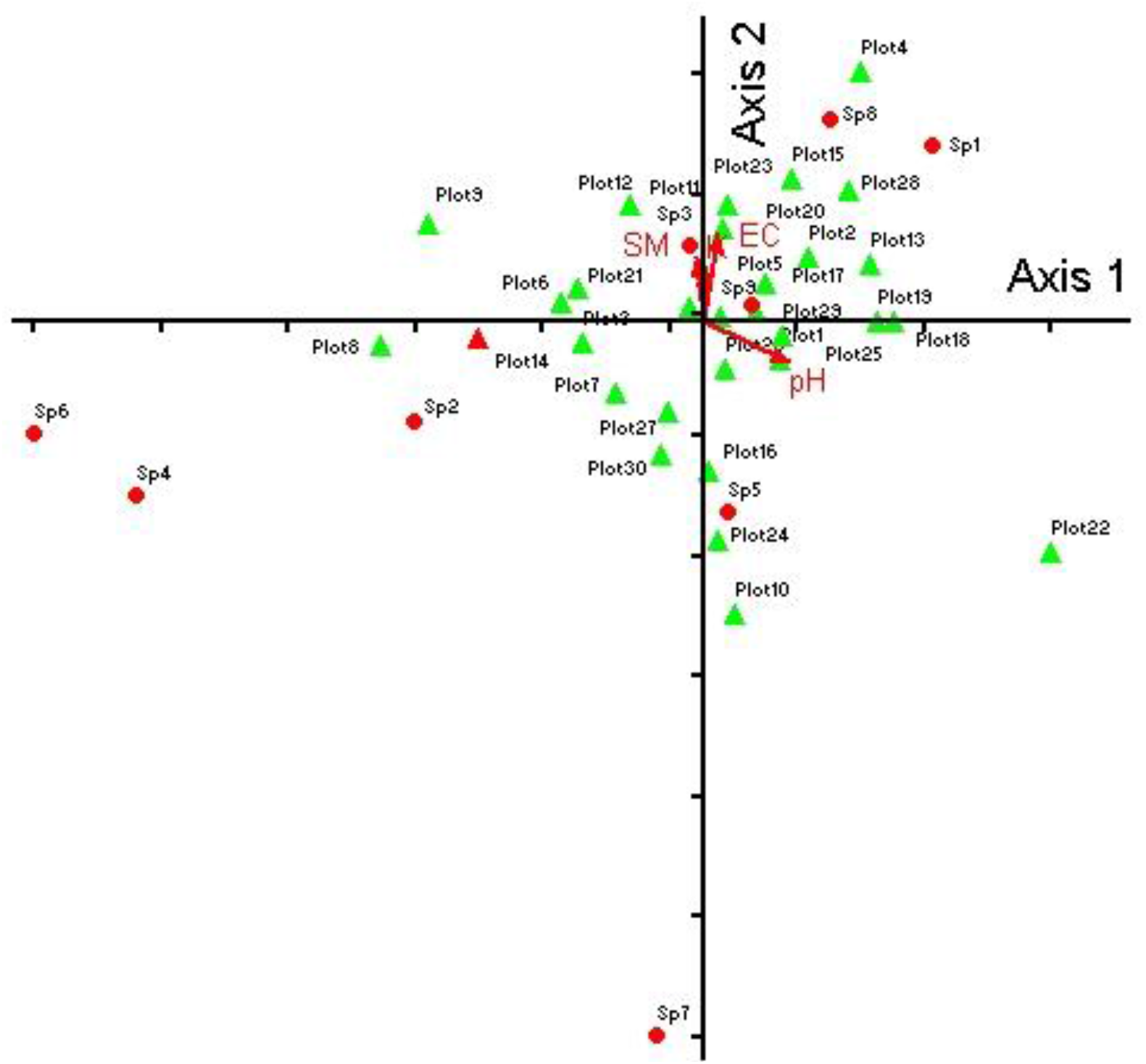
Graph showing the correlation between soil parameters and *Rhododendron kesangiae*

The species composition and electrical conductivity (EC) exhibited moderate to significant positive associations (*r* = 0.042 - 0.794, *r-sq* = 0.002 - 0.631, *tau* = 0.175 - 0.621). Temperature showed weak to moderate correlations (*r* = -0.176 - 0.304, *r-sq* = 0.031 - 0.092, *tau* = 0.106 - 0.161), whereas Slope had weak correlations (*r* = -0.019 - 0.132, *r-sq* = 0.000 - 0.017, *tau* = 0.018 - 0.161). The species composition correlation with precipitation (Ppt) and altitude (Alt) showed weak to moderate relationships (*r* = -0.249 - 0.314, *r-sq* = 0.062 - 0.099, *tau* = 0.200 - 0.223 and *r* = -0.171 - 0.293, *r-sq* = 0.029 - 0.086, *tau* = 0.074 - 0.175), respectively.

### 3.6 Soil components of *Rhododendron kesangiae*

Important information about the ecological factors influencing *Rhododendron kesangiae* growth and regeneration are provided by Pearson’s correlation analysis between the plant’s regeneration and several soil parameters (soil moisture, SOC, SN, SOM, SP, SK, pH, EC). Soil parameters such as soil moisture, soil NPK, pH and electrical conductivity were tested for *Rhododendron kesangiae*. The average soil pH was 5.18, with a range of 4.7 to 6.09, indicating that *Rhododendron kesangiae* requires slightly acidic to neutral soil for its proper growth and regeneration. On average, soil MC was 28.8%, SOC was 1.38, N was 0.11, SOM was 2.3, P was 0.64, and K was 421.1, indicating that it requires well-moist soil and high levels of soil nutrients for its proper growth and development.

**Table 3:**
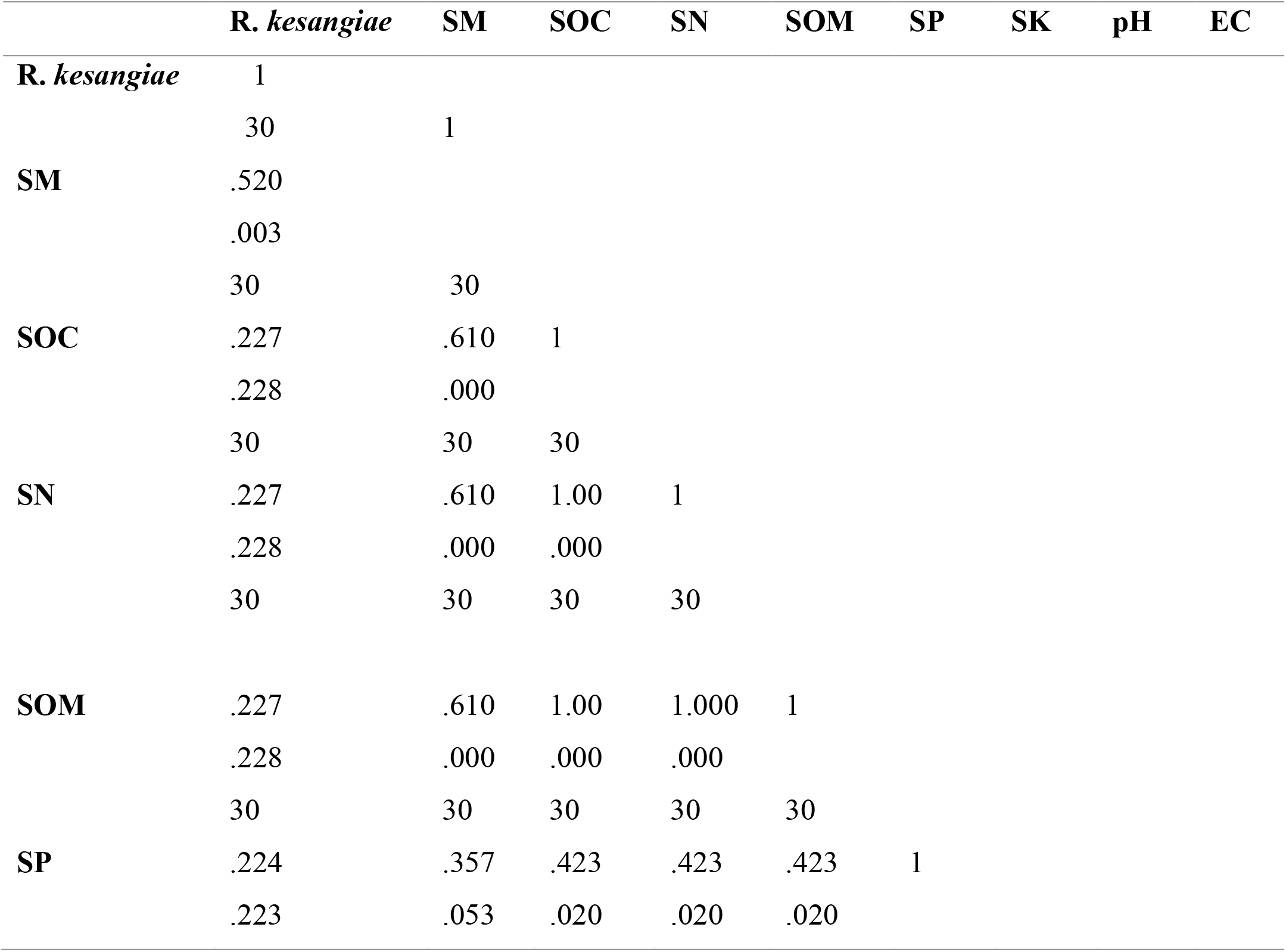
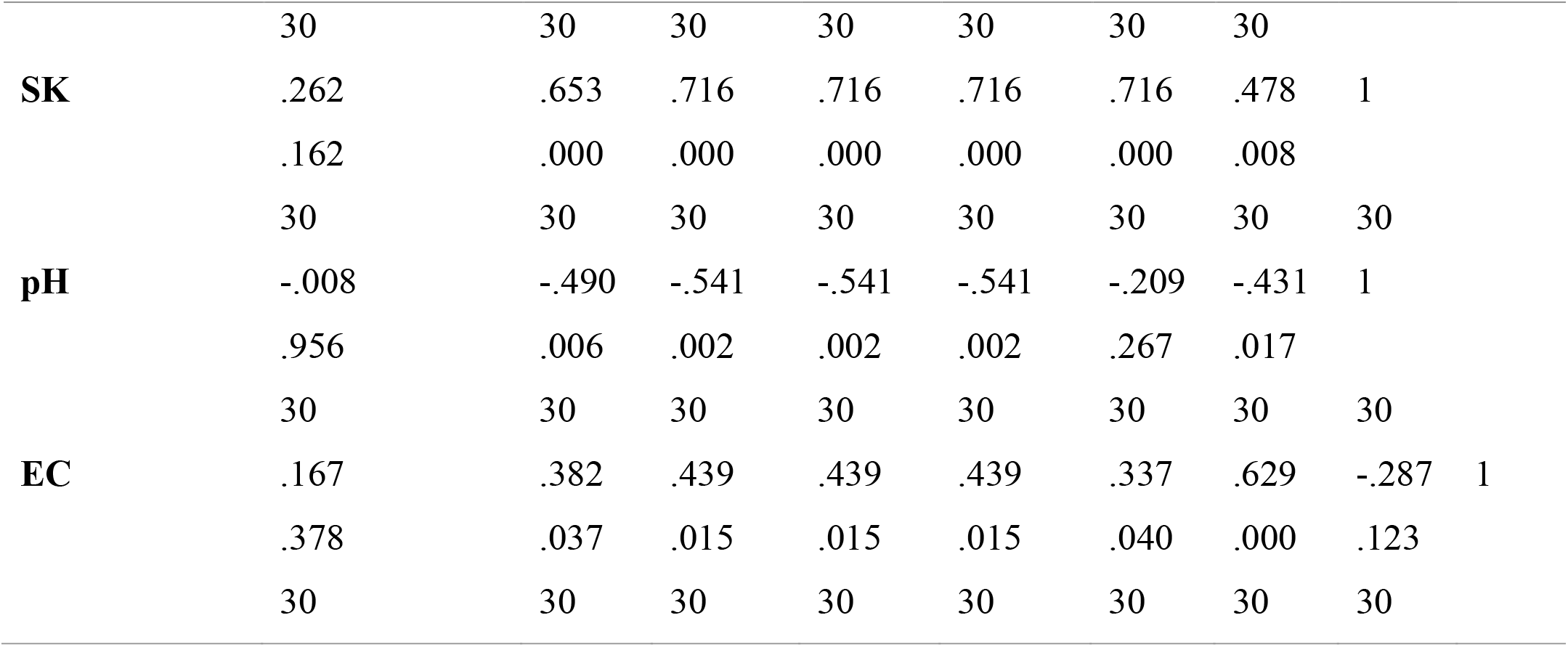
Pearson’s correlation between R. *kesangiae* and soil parameter.

Pearson’s correlation analysis was conducted to see the association between *Rhododendron kesangiae* regeneration and different soil parameters. The findings showed multiple significant associations between soil parameters and *Rhododendron kesangiae* regeneration. There was a significant positive association between soil moisture (*r* = 0.520, *p* = 0.003).

However, the plant’s regeneration had a negative correlation with soil pH (*r* = -0.008). These results show that soil phosphorus, potassium, organic carbon, and moisture may all affect the growth and regeneration of *Rhododendron kesangiae*.

### 3.7 Relation between *Rhododendron kesangiae* and environmental variables

This study examined the association or relationship between *Rhododendron kesangiae* regeneration and various environmental factors such as slope, temperature, rainfall and altitude using correlation analysis. The result of the analysis showed a strong positive relationship between regeneration and precipitation (*r* = 0.371), indicating that higher rainfall leads to better R. *kesangiae* regeneration.

**Table 4:**
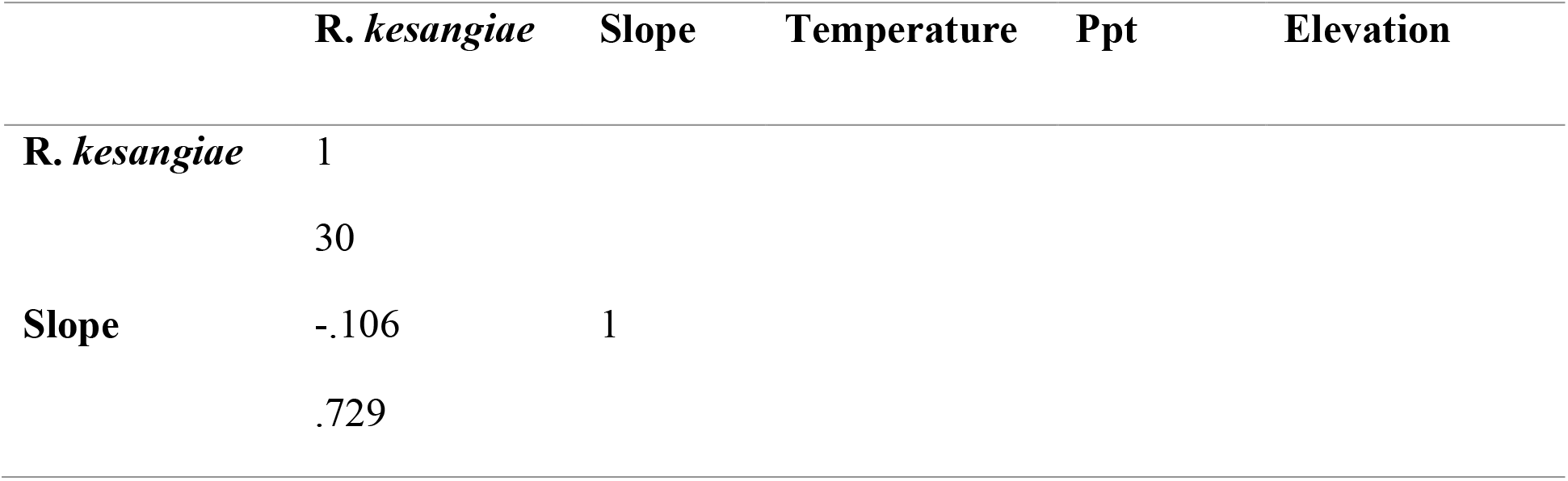
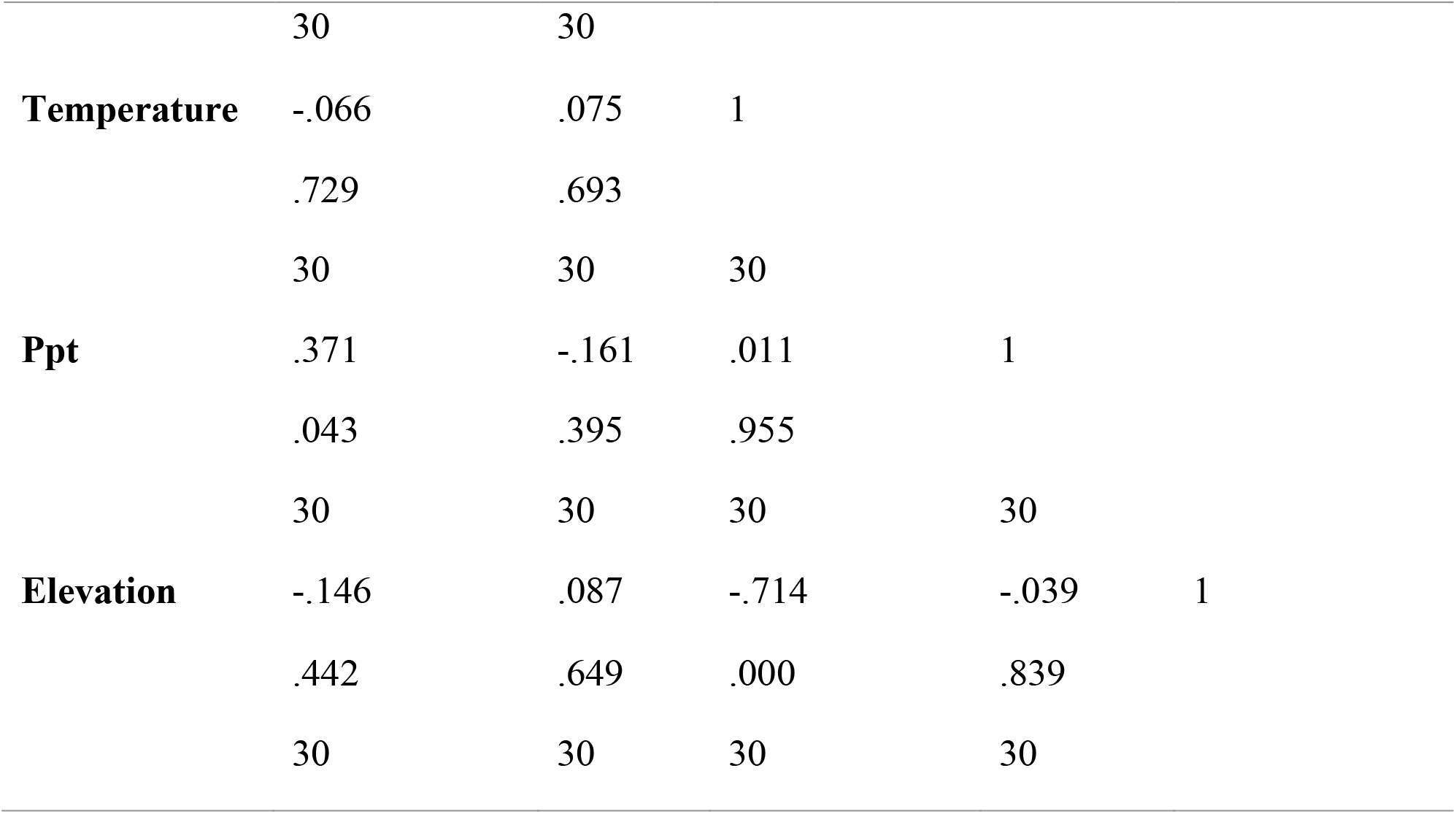
Pearson’s correlation between R. *kesangiae* and environmental variables.

|  | **R. kesangiae** | **Slope** | **Temperature** | **Ppt** | **Elevation** |
| --- | --- | --- | --- | --- | --- |
| **R. kesangiae** | 1 |  |  |  |  |
|  | 30 |  |  |  |  |
| **Slope** | -.106 | 1 |  |  |  |
|  | .729 |  |  |  |  |
|  | 30 | 30 |  |  |  |
| **Temperature** | -.066 | .075 | 1 |  |  |
|  | .729 | .693 |  |  |  |
|  | 30 | 30 | 30 |  |  |
| **Ppt** | .371 | -.161 | .011 | 1 |  |
|  | .043 | .395 | .955 |  |  |
|  | 30 | 30 | 30 | 30 |  |
| **Elevation** | -.146 | .087 | -.714 | -.039 | 1 |
|  | .442 | .649 | .000 | .839 |  |
|  | 30 | 30 | 30 | 30 |  |

The regeneration of *Rhododendron kesangiae* and altitude had a negative correlation regeneration and altitude (*r* = -0.146), indicating that the plant regenerates better at lower altitudes. The overall findings suggest that precipitation may have a significant impact on R. *kesangiae* (*p* = 0.043) regeneration, while slope, temperature, and elevation do not seem to have a significant impact on its regeneration. The correlation analysis between R. *kesangiae* regeneration and several environmental factors fulfils the objective of understanding the ecological parameters that influence the growth and regeneration of *Rhododendron kesangiae*.

### 3.8 Conservation threats to *Rhododendron kesangiae*

The primary threats to *Rhododendron kesangiae* include habitat loss, fragmentation, pollution, tourism, cattle raising, and climate change. We utilised Miradi software to identify the likely causes of each risk and describe conservation goals and solutions. Conservation threats such as habitat degradation and fragmentation can threaten the survival and well-being of the species. Habitat degradation and fragmentation due to habitat loss, anthropogenic activities, infrastructure/urban development, and land conversions can lead to a decline in suitable habitats for *Rhododendron kesangiae* and can disrupt gene flow leading to local extinction (lbanez et al., 2014).

Tourism is also identified as a threat to *Rhododendron kesangiae* due to the increasing popularity of the Lungchutse trail among tourists, which runs through the natural habitat of *Rhododendron kesangiae*. Tourism activities can introduce pollutants, generate waste, and disturb the habitat of the species.

It was observed in the Babia Gora National Park that tourists caused damage to plant communities by trampling, causing mechanical damage, erosion of trail surface and generating waste (Lamorski & Dabrowski, 2010).

**Figure 5:**
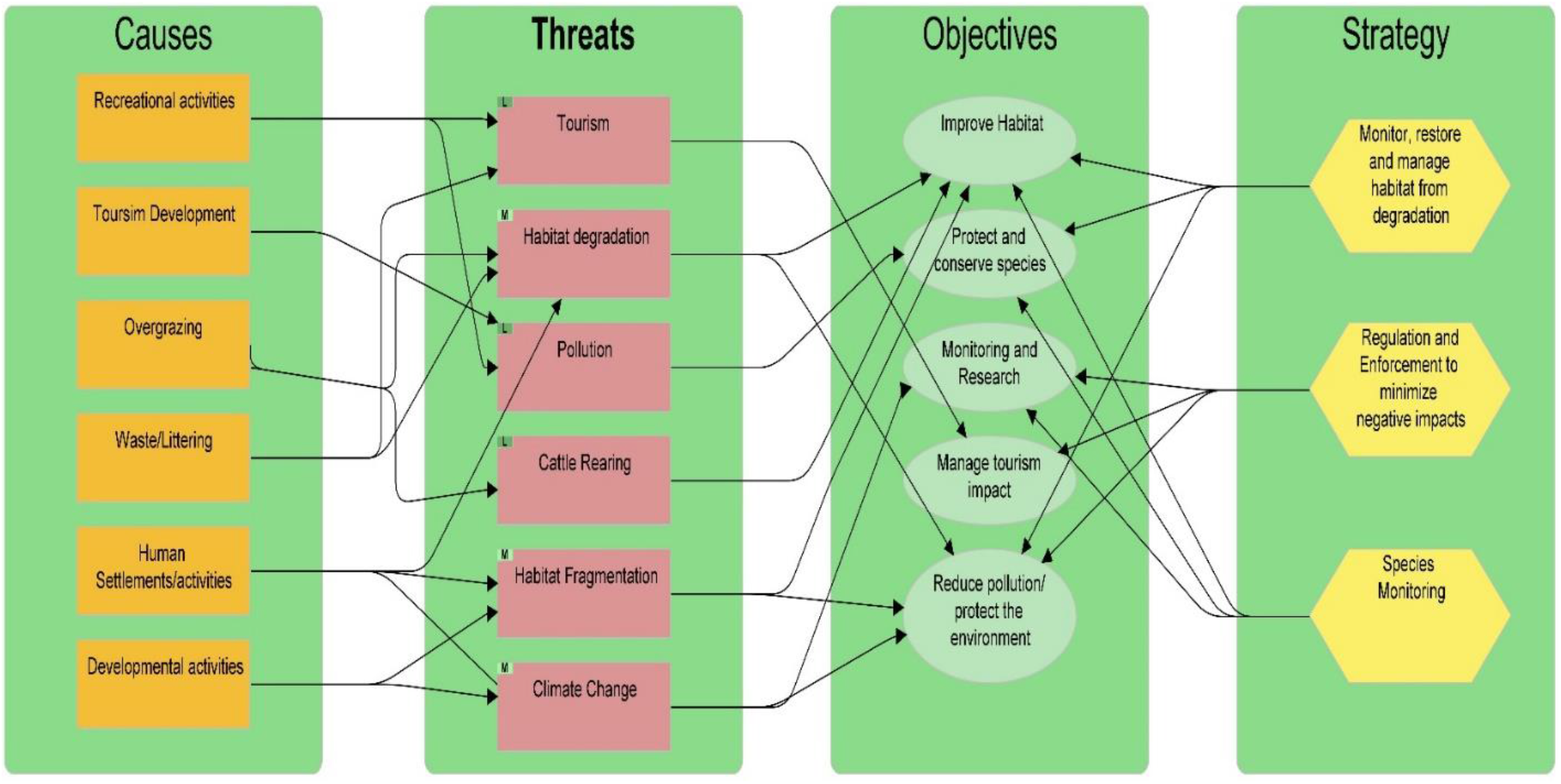
Threat analysis of R. *kesangiae* using Miradi

Cattle rearing is also identified as a threat to *Rhododendron kesangiae* as observed in the field; grazing was a prominent situation in most of the plots as determined by the cattle droppings. Cattle can trample the plants and browse on young regenerating shoots. Climate change can have a significant effect on *Rhododendron kesangiae* by altering its spatial distribution. The changes in temperature and precipitation patterns can shift the habitat of R. kesangiae and lead to changes in its distribution, which is already limited to certain areas. The change in climatic conditions can also influence flowering and reproduction in plants such as R. *kesangiae*, which may affect their pollination, reproductive success, and growth.

To address these threats, conservation strategies can be developed, such as habitat restoration through reforestation, proper pollution control, livestock management (fencing off sensitive areas), and climate mitigation programs such as carbon sequestration.

Aligned with objectives that are specific, achievable, measurable, relevant and time-bound (SMART), such as habitat improvement, monitoring and reducing pollution, eco-friendly tourism, and conducting research.

Through this Miradi analysis, it can also be drawn that *Rhododendron kesangiae* is on the verge of being vulnerable due to various conservation threats it is facing in its natural habitats. The findings of this Miradi analysis contribute to the objective of identifying conservation threats to *Rhododendron kesangiae* by providing an understanding of the threats to the species in its natural habitat and guiding conservation actions to ensure its long-term existence in the wild. Implementing our conceptual model provides a strategic and structured approach to ensure *Rhododendron kesangiae* is in its natural habitat. Understanding the threats, addressing causes, and setting objectives can help preserve and protect the species in the wild.

Although the species is listed as least concerned, a species listed as least concerned may face new challenges tomorrow, which can lead to the decline of the species. Proactive studies such as identifying the conservation threat to the species by using Miradi can act as an early warning system to identify and mitigate the threats to the species to prevent possible future decline of the species. It also contributes to the global diversity goals and targets as outlined in the Convention on Biological Diversity (CBD), which states that even the least concerned species should be monitored and protected to maintain ecological balance.

## 4. Conclusion and Recommendation

This study was conducted with the objective of exploring the regeneration and conservation status of *Rhododendron kesangiae*, a species endemic to Bhutan. The study investigated the current state of regeneration of R. *kesangiae* and its relationship with several environmental factors such as slope, rainfall, elevation, temperature, and soil characteristics in the study area. Conservation risks to the growth and survival of the species were also identified to ensure its long-term survival in the wild.

The study found that the current state of regeneration of Rhododendron *kesangiae* was fair, with more seedlings and saplings than mature individuals (141 seedlings ≤ 184 saplings > 129 adults) in the study area. This finding indicates and assures conservationists that the likelihood of the species’ population survival may increase with appropriate conservation efforts in place. Understanding the environmental factors that influence the regeneration of the species, such as soil moisture, soil nutrients (NPK) and precipitation, gives information for preserving the required environmental conditions. The plant’s association with other tree species, such as *Tsuga dumosa, Betula utilis*, and *Abies densa*, suggests that forest communities need to be protected to ensure their long-term survival and understand how these species interact with each other. The threat assessment for R. *kesangiae* regeneration generated using Miradi can support and guide conservation measures, providing stakeholders and policymakers with a clear path for targeted interventions to ensure *Rhododendron kesangiae’s* survival in its natural habitat.

While this study focuses on the regeneration ecology and conservation threats of the species in its natural habitat, researchers who are interested in this field of study could explore *Rhododendron kesangiae* on a genetic level and explore its microhabitat preferences.

## Acknowledgment

We acknowledge the Department of Forest and Park Services, Bhutan, for allowing us to carry out the research in the National Botanical Park in Lungchutse. We also acknowledge the College of Natural Resources for providing necessary logistic and technical support during the research time.

## Conflict of Interest

The authors declare no conflict of interest.

## Author contribution statement

Study conception and design by **CTD & YD**; Methodology Implementation by **CTD & YD**; Data collection by **KTR**; Data analysis/Interpretation by **CTD & YD**; Manuscript writing/ revision by **CTD & YD**. All authors gave final approval.

